# Cryo-EM structures of Tau filaments from the brains of mice transgenic for human mutant P301S Tau

**DOI:** 10.1101/2023.08.31.555656

**Authors:** Manuel Schweighauser, Alexey G. Murzin, Jennifer Macdonald, Isabelle L. Lavenir, R. Anthony Crowther, Sjors H.W. Scheres, Michel Goedert

## Abstract

Mice transgenic for human mutant P301S tau are widely used as models for human tauopathies. They develop neurodegeneration and abundant filamentous inclusions made of human mutant four-repeat tau. Here we used electron cryo-microscopy (cryo-EM) to determine the structures of tau filaments from the brains of Tg2541 and PS19 mice. Both lines express human P301S tau (0N4R for Tg2541 and 1N4R for PS19) on mixed genetic backgrounds and downstream of different promoters (murine *Thy1* for Tg2541 and murine *Prnp* for PS19). The structures of tau filaments from Tg2541 and PS19 mice differ from each other and those of tau filaments from human brains. However, they share a substructure at the junction of repeats 2 and 3, which comprises residues I297-V312 of tau and includes the P301S mutation. The filament core from the brainstem of Tg2541 mice consists of residues K274-H329 of tau and two disconnected protein densities. Two non-proteinaceous densities are also in evidence. The filament core from the cerebral cortex of line PS19 extends from residues G271-P364 of tau. One strong non-proteinaceous density is also present. Unlike the tau filaments from human brains, the sequences following repeat 4 are missing from the cores of tau filaments from the brains of Tg2541 and PS19 mice.

## INTRODUCTION

Identification of dominantly inherited mutations in human *MAPT*, the tau gene, established that dysfunction of tau protein is sufficient to cause neurodegeneration and dementia [1–3]. To date, 67 different mutations in *MAPT* are known to cause frontotemporal dementia and parkinsonism linked to chromosome 17 (FTDP-17T) [4]. They are gene dosage, exonic (missense and deletion) and intronic mutations. FTDP-17T brains exhibit atrophy of the frontal and temporal lobes of the neocortex, striatum and substantia nigra, with accompanying nerve cell loss and gliosis. Abundant argyrophilic cytoplasmic inclusions containing filamentous tau are present in nerve cells and, in some cases, glia.

In the brains of human adults, six tau isoforms are expressed from a single gene through alternative mRNA splicing [5]. They differ by the presence (1N, 2N) or absence (0N) of two inserts in the N-terminal half and an insert in the C-terminal half. The latter consists of a repeat of 31 amino acids (encoded by exon 10), giving rise to three isoforms with four repeats (4R). The other three isoforms have three repeats (3R). Exon 10 encodes R2 of the 4R tau isoforms. The repeats and some adjoining sequences constitute the microtubule-binding domains of tau [6]. They also form the cores of assembled tau in neurodegenerative diseases, suggesting that physiological function and pathological assembly are mutually exclusive.

Besides their conceptual importance, mutations in *MAPT* have made it possible to produce and characterise transgenic mouse lines that exhibit tau hyperphosphorylation, filament assembly and neurodegeneration. Mouse lines transgenic for human P301L or P301S tau have been the most widely studied [7–10]. In humans, mutation P301S tau causes an early-onset form of FTDP-17T [11–16] It reduces the ability of tau to interact with microtubules [11] and promotes heparin-induced filament assembly of tau [17], without influencing the splicing of exon 10 of *MAPT* [18].

Here we report the electron cryo-microscopy (cryo-EM) structures of tau filaments from the brains of Tg2541 and PS19 mice transgenic for human P301S tau [9,10]. They express 0N4R tau and 1N4R tau, respectively. The filament structures are unlike those of inclusions from human brains. They are also different between the two mouse lines, which used distinct promoters (murine *Thy1* in [9] and murine *Prnp* in [10]).

## MATERIALS AND METHODS

### Transgenic mice

Animal experiments were conducted in accordance with the UK Animals (Scientific Procedures) Act of 1986, with local ethical approval (MRC Laboratory of Molecular Biology Animal Welfare and Ethical Review Body).

Tg2541 mice express full-length human tau (0N4R) with the P301S mutation under the control of the murine *Thy1* promoter on a mixed C57BL/6 x CBA background [9]; homozygous mice develop amyloid inclusions made of hyperphosphorylated mutant 4R tau in the central nervous system from approximately 2 months of age [9, 19–21]. PS19 mice express full-length human tau (1N4R) with the P301S mutation under the control of the murine *Prnp* promoter on a mixed C57BL/6 x C3H background [10]; heterozygous mice develop filamentous inclusions made of mutant 4R tau in the central nervous system from approximately 6 months of age. Breeding pairs of PS19 mice were obtained from the Jackson Laboratory.

### Filament extraction

Sarkosyl-insoluble material was extracted from the brainstem and spinal cord of homozygous Tg2541 mice ranging from 4 weeks to 25 weeks of age and from the cerebral cortex of heterozygous 48-week-old PS19 mice, as described [22]. Tissues were homogenised in 20 vol buffer A (10 mM Tris-HCl, pH 7.5, 0.8 M NaCl, 10% sucrose and 1 mM EGTA), brought to 2% sarkosyl and incubated for 30 min at 37° C. The samples were centrifuged at 10,000 g for 10 min, followed by spinning of the supernatants at 100,000 g for 25 min. The pellets were resuspended in 700 μl/g extraction buffer and centrifuged at 5,000 g for 5 min. The supernatants were diluted threefold in 50 mM Tris-HCl, pH 7.4, containing 0.15 M NaCl, 10% sucrose and 0.2% sarkosyl, and spun at 166,000 g for 30 min. The pellets were resuspended in 50 μl/g 20 mM Tris-HCl, pH 7.4, 100 mM NaCl.

### Immunogold negative stain electron microscopy

Immunogold negative stain electron microscopy was carried out as described [23]. Anti-tau antibodies BR133, BR136, anti-4R, BR135, TauC4 and BR134 were used at 1:50. BR136 [24], anti-4R [25], BR135 [5] and TauC4 [26] are specific for R1, R2, R3 or R4 of tau. BR133 and BR134 are specific for the amino-or carboxy-ends of tau, respectively [5].

### Electron cryo-microscopy

Three μl of the sarkosyl-insoluble fractions were applied to glow-discharged (Edwards S150B) holey carbon grids (Quantifoil AuR1.2/1.3, 300 mesh) that were plunge-frozen in liquid ethane using a Vitrobot Mark IV (Thermo Fisher Scientific) at 100% humidity and 4° C. Cryo-EM images were acquired using EPU software on a Titan Krios microscope (Thermo Fisher Scientific) operated at 300 kV. For Tg2541 mice aged 24 weeks, movies were acquired on a Gatan K2 Summit detector using a pixel size of 1.15 Å. For Tg2541 mice aged 8 weeks, images were acquired on a Gatan K3 detector using a pixel size of 0.93 Å. For both detectors, a quantum energy filter with a slit width of 20 eV was used to remove inelastically scattered electrons. For PS19 mice aged 48 weeks, movies were acquired on a Falcon-4 detector at a pixel size of 0.824 Å. See Supplementary Table for further details.

### Helical reconstruction

Datasets were processed in RELION using standard helical reconstruction [27]. Movie frames were gain-corrected, aligned and dose-weighted using RELION’s own motion correction programme [28]. Contrast transfer function (CTF) was estimated using CTFFIND4-1 [29]. Filaments were picked manually. Three-dimensional auto-refinements were performed with optimisation of the helical twist and rise parameters once the resolutions extended beyond 4.7 Å. To improve the resolution, Bayesian polishing and CTF refinement were used [30]. Final maps were sharpened using standard post-processing procedures in RELION and resolution estimates calculated based on the Fourier shell correlation (FSC) between two independently refined half-maps at 0.143 [31] (Supplementary Figure 1).

### Model building and refinement

The atomic models were docked manually in the densities using Coot [32]. Model refinements were performed using *Servalcat* [33] and REFMAC5 [34,35]. Models were validated with MolProbity [36]. Figures were prepared with ChimeraX [37] and PyMOL [38].

## RESULTS

### Time course of tau filament formation in Tg2541 mice

Sarkosyl-insoluble material was extracted from the brainstem of Tg2541 mice aged 4, 8, 12, 16 and 24 weeks. Filaments were observed at 4 and 8 weeks, with a progressive increase until 24 weeks (Figure 1a). In the spinal cord, some filaments were in evidence at 5, 6 and 8 weeks, with a marked increase at 12, 16 and 25 weeks. They were decorated by anti-tau antibody BR134 (Figure 1b). When analysed at 20 weeks of age, filaments from cerebral cortex, brainstem and spinal cord were decorated by antibodies BR133, TauC4 and BR134, but not by BR136, anti-4R or BR135 (Figure 2).

**Figure 1.**
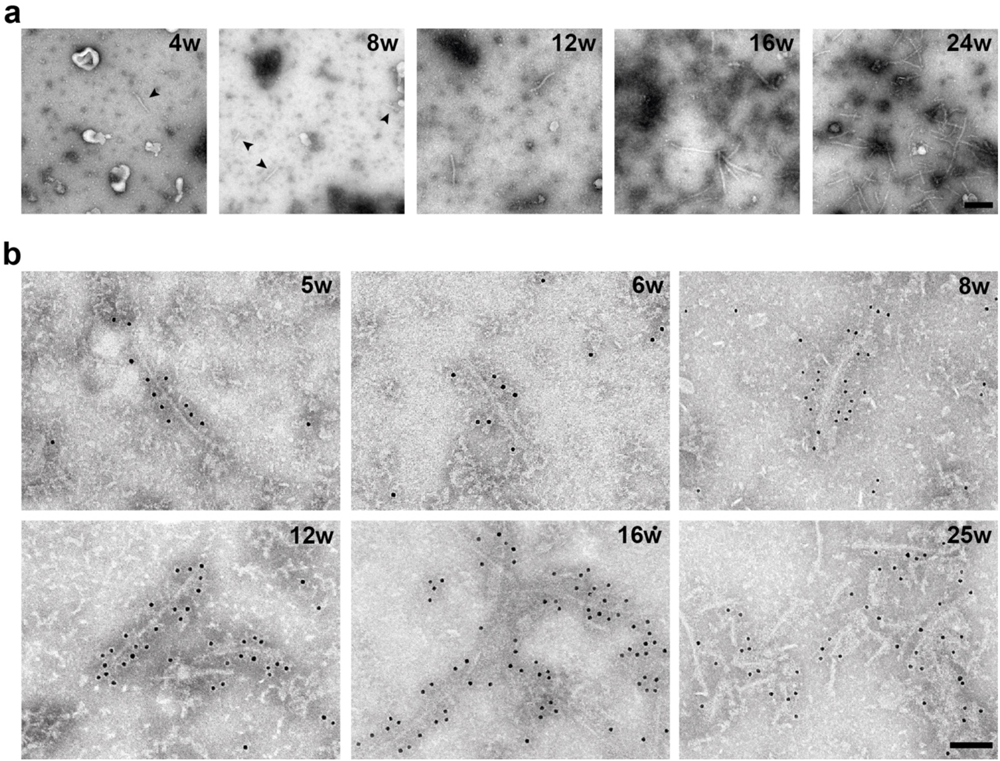
Tau filaments from the central nervous system of Tg2541 mice aged 4-25 weeks. **a,** Negative-stain electron microscopy images of filaments extracted from the brainstem of Tg2541 mice aged 4-24 weeks (w). Arrowheads in the first two panels point to short filaments. Scale bar, 200 nm. **b,** Immunogold negative-stain electron microscopy of Tau filaments extracted from the spinal cord of Tg2541 mice aged 5-25 weeks (w). Anti-tau antibody BR134 was used. Note the gold particles lining the filament surfaces. Scale bar, 100 nm.

**Figure 2.**
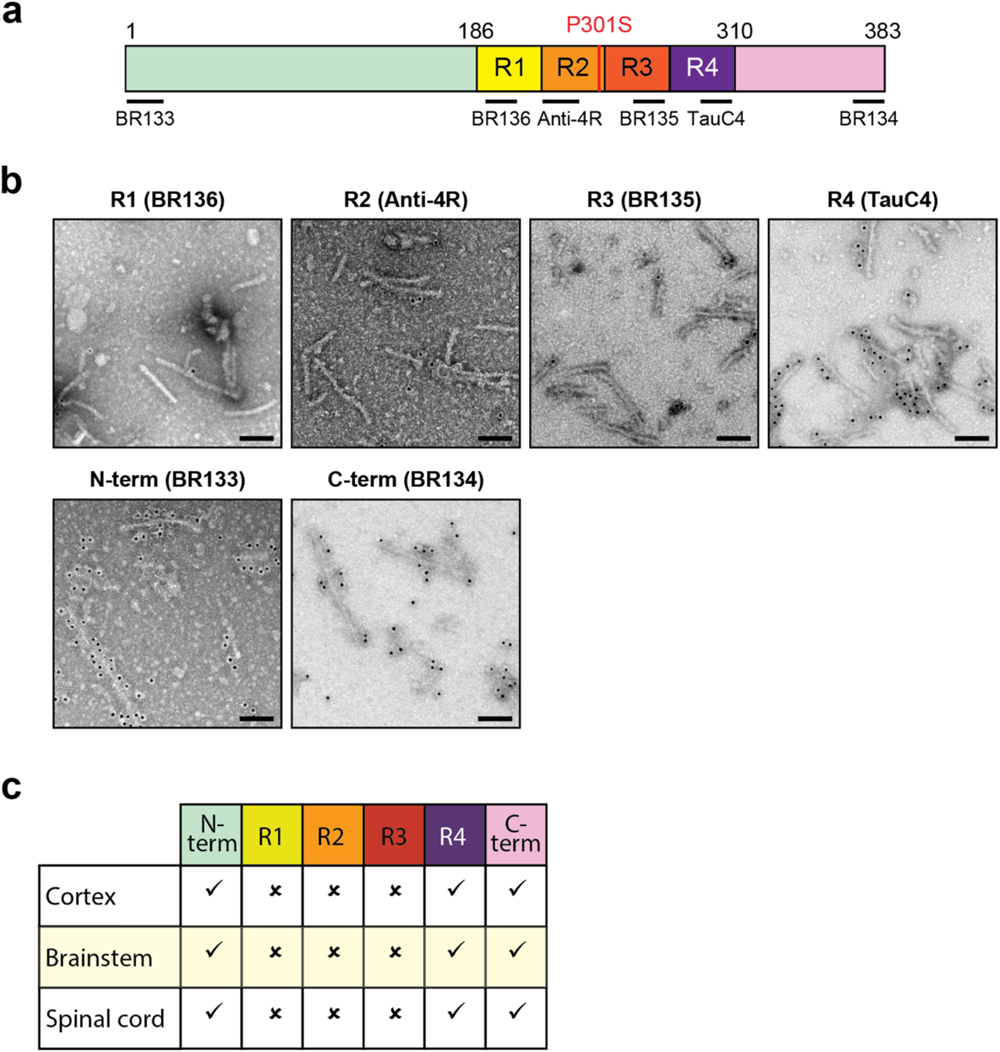
Immunogold negative-stain electron microscopy of Tau filaments extracted from the central nervous system of 24-week-old Tg2541 mice. **a,** Diagram of 0N4R P301S tau showing the repeats (R1-R4) and the epitopes of antibodies BR133 (N-terminus), BR136 (R1), anti-4R (R2), BR135 (R3), TauC4 (R4) and BR134 (C-terminus). **b,** Immunogold negative-stain electron microscopy of Tau filaments from the brainstem with BR136, anti-4R, BR135, TauC4, BR133 and BR134. **c,** Table summarising results from immunogold negative-stain electron microscopy of Tau filaments from frontal cortex, brainstem and spinal cord. Tick marks indicate antibody decoration of filaments and crosses indicate that the antibodies did not decorate filaments. Filaments were decorated by BR133, TauC4 and BR134, but not by BR136, anti-4R and BR135. Scale bar, 100 nm.

### Cryo-EM structures of tau filaments from Tg2541 mice

Cryo-EM of tau filaments from line Tg2541 extracted at 8 weeks and 24 weeks revealed the same structure (Figure 3). The filament core comprises a contiguous segment, spanning residues K274-H329, and two disconnected protein densities of about 11 residues each, the sequences of which could not be assigned unambiguously. The contiguous segment, consisting of the last residue of R1, the whole of R2 and most of R3, forms two layers. One layer is formed by the whole of R2 and contains strands β1-β4, whereas the second layer consists of the residues of R3 that are arranged in strands β5-β7. The turn between layers is filled by hydrophobic residues I297 and V300 from R2, and V306, I308 and Y310 from R3; it is further stabilised by a salt bridge between D295 and K311.

**Figure 3.**
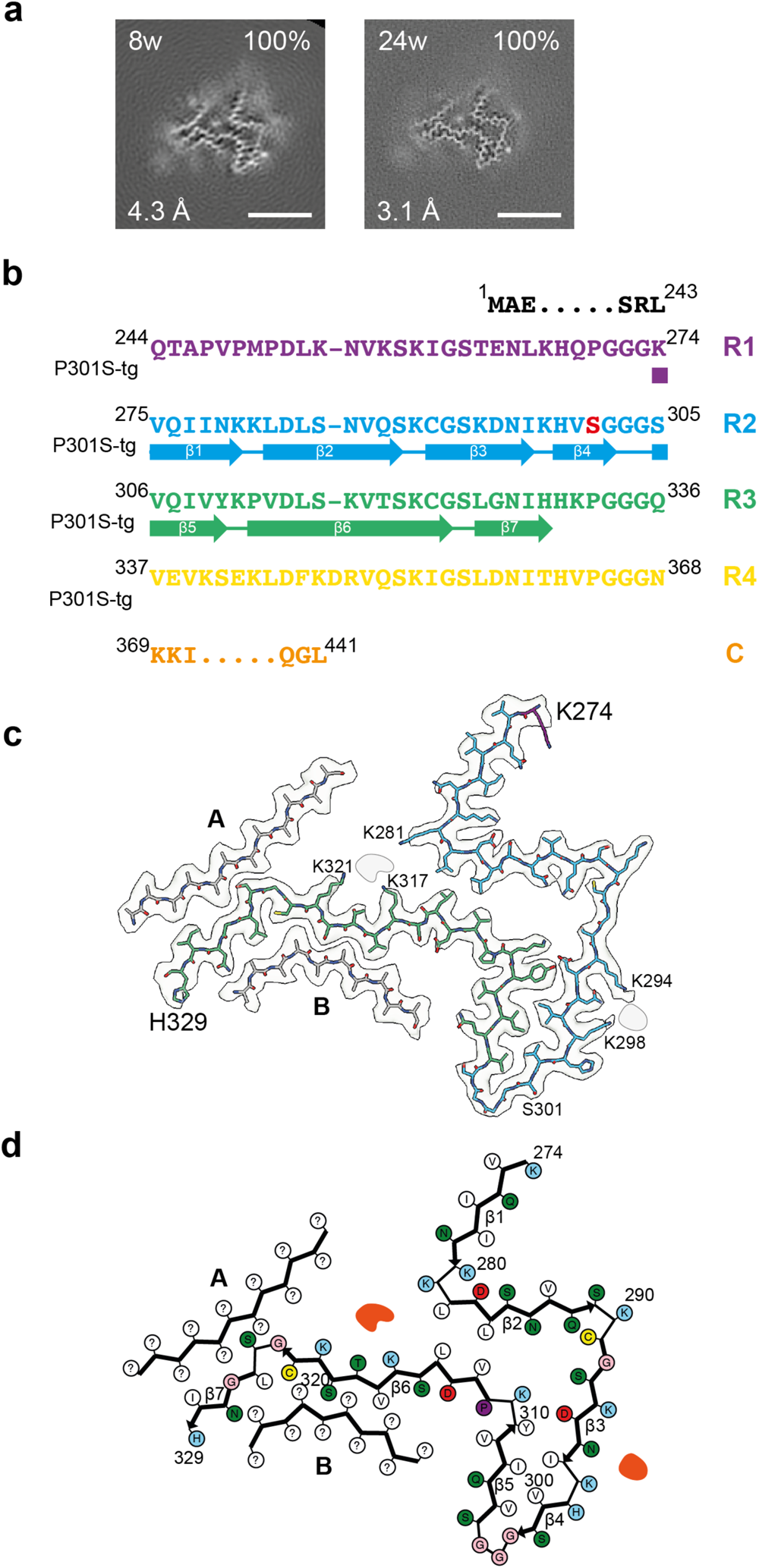
Structure of P301S Tau filaments from the brainstem of Tg2541 mice. **a,** Cross-sections through the cryo-EM reconstructions, perpendicular to the helical axis and with a projected thickness of approximately one rung, are shown for the P301S Tau filaments from the brainstem of Tg2541 mice aged 8 weeks and 24 weeks. The cryo-EM map resolution (bottom left) and the percentages of a given filament type among the filaments in the datasets (top right) are shown. Scale bars, 5 nm. **b,** Sequence alignment of the microtubule-binding repeats (R1-R4) of Tau with the observed seven ý-strand regions (ý1-ý7, arrows). Mutant serine residue at position 301 is highlighted in red. **c,** Sharpened high-resolution cryo-EM map of the P301S Tau filaments with the atomic model overlaid. Residues in R1-R4 and the C-terminal domain are coloured purple, blue, green, yellow and orange, respectively. Islands A and B are indicated in grey. **d,** Schematic of the Tg2541 Tau filament fold. Negatively charged residues are shown in red, positively charged residues in blue, polar residues in green, non-polar residues in white, sulphur-containing residues in yellow, prolines in purple and glycines in pink. Thick connecting lines with arrowheads indicate ý-strands (ý1-ý7). Additional densities are shown in red. Unknown residues are indicated by question marks.

The serine side chain of the P301S mutation site sits on the outside surface of this turn. The middle parts of both layers are held together by hydrophobic interactions between residues L282 and L284 from R2 and V313 and L315 from R3. On the turn side of this hydrophobic cluster, there is a solvent-filled cavity that is inlaid with the polar side chains of N286, Q288, S293, D295 and K311, whereas on the other side both layers diverge and point in opposite directions. The N-terminal strand β1 of the R2 layer makes only few interactions with other protein regions, whereas the C-terminal part of the R3 layer is covered on both sides by disconnected density islands A and B.

Both islands lack side chain densities beyond Cβ atoms, preventing unambiguous sequence assignment. However, some clues as to their identity are provided by proximity to the C-terminal part of layer R3 at certain positions. Thus, the residue in island A that packs against the turn ^324^GS^325^ must have a small side chain, as does the residue in island B that packs between V318 and S320. The nearest candidate for either ‘anchor’ position is S341 in tau. Modelling the sequence around this residue into the islands’ densities shows that both islands can accommodate well R4 segment ^336^QVEVKSEKLDF^346^, making favourable interactions in their interfaces with layer R3. In contrast, for the alternative assignment of island A with the ‘nearest’ R1 segment, ^253^LKNVKSKIGST^263^, residue S258, which is the first small residue N-terminal to K274 that can reach the ‘anchor’ point, results in a buried position of the charged group of K254 without compensation and places G261 in a site occupied by a residue with a side chain.

The same R4 segment cannot fill both islands to full occupancy on its own. If it alternates between islands from one rung to the next, other segments are needed to fill the gaps. This role can be taken by the next R4 segment ^347^KDRVQSKIGSL^357^, which runs anti-parallel to the first, with S352 occupying the ‘anchor’ position in both islands. Modelling the ý-hairpins made of both segments with the turn at ^346^FK^347^ into the densities of both islands shows that they fit well (Supplementary Figure 2). This suggests that the core sequence of tau filaments from the brainstem of Tg2541 mice extends up to L357 and includes most of R4.

The proposed β-hairpin islands would be connected to the contiguous segment via a six-residue turn ^330^HKPGGG^335^ that is expected to be less ordered, since its position relative to the ordered core will alternate between rungs.

Apart from the protein density islands, there are two non-proteinaceous densities, which correspond to putative cofactors and/or post-translational modifications. One density resides on the side of the R2 layer and is co-ordinated by K294 and K298, whereas the other density sits at the divergence point of layers R2 and R3, and is co-ordinated by K281, K317 and K321. Similar non-proteinaceous densities that are co-ordinated by lysine residues have been observed in 4R tau filaments extracted from *post mortem* human brains [39,40].

### Cryo-EM structures of tau filaments from PS19 mice

The filament core from the cerebral cortex of 48-week-old mice from line PS19 extends from residues G271-P364 of tau and consists of the C-terminal four residues of R1, the whole of R2 and R3 and 28 N-terminal residues of R4 (Figure 4). Its secondary structure comprises eight ý-strands that range from 3-23 residues in length. They form a hairpin-like structure, the tip of which folds back at the turns between β2 and β3, and between β4 and β5, giving the cross-section of the filament core a tadpole-like shape.

**Figure 4.**
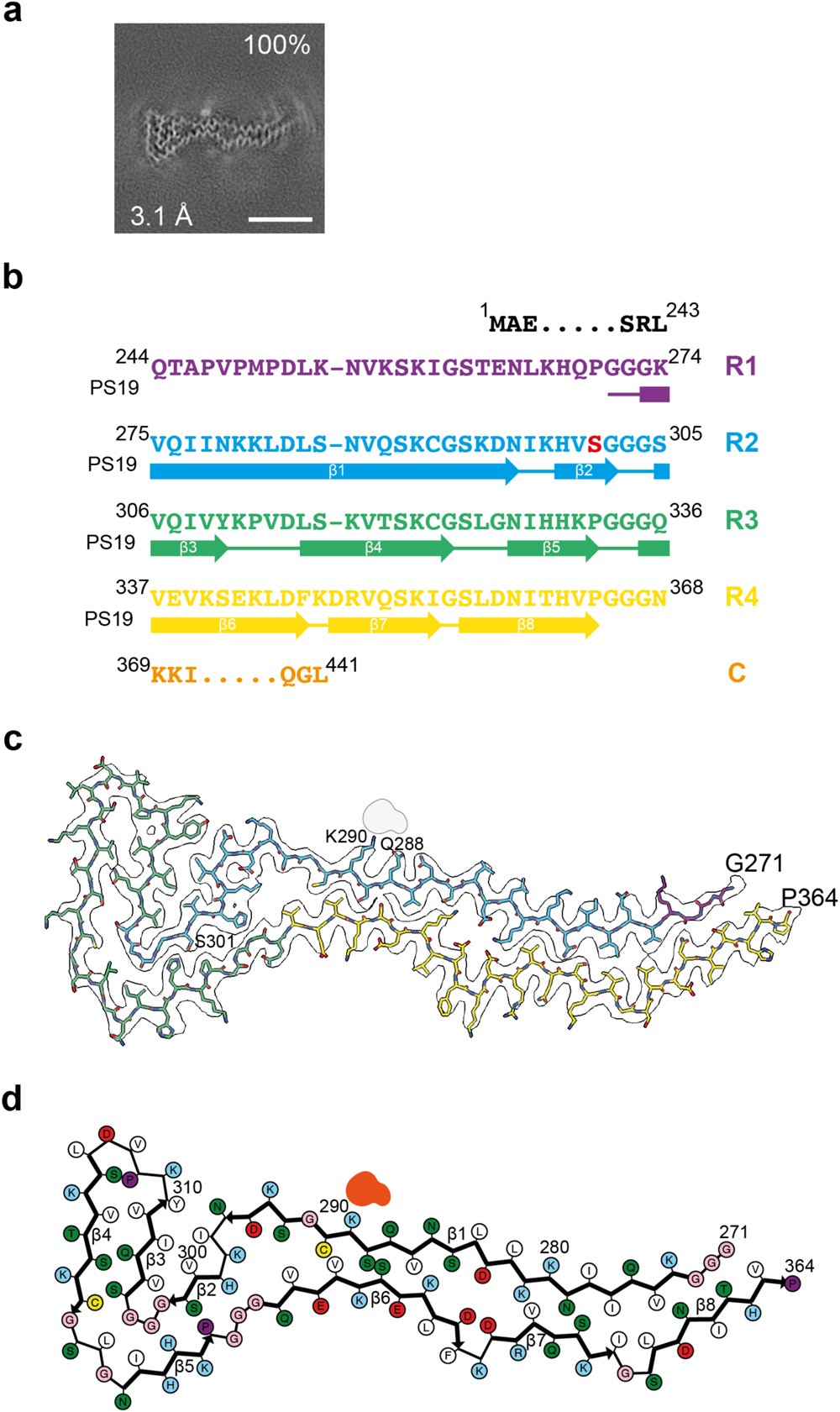
Structure of P301S Tau filaments from the cerebral cortex of PS19 mice. **a,** Cross-section through the cryo-EM reconstructions, perpendicular to the helical axis and with a projected thickness of approximately one rung, are shown for the P301S Tau filaments from the cerebral cortex of PS19 mice aged 48 weeks. The cryo-EM map resolution (bottom left) and the percentages of a given filament type among the filaments in the datasets (top right) are shown. Scale bar, 5 nm. **b,** Sequence alignment of the microtubule-binding repeats (R1-R4) of Tau with the observed eight ý-strand regions (ý1-ý8, arrows). Mutant serine residue at position 301 is highlighted in red. **c,** Sharpened high-resolution cryo-EM map of the P301S Tau filaments with the atomic model overlaid. Residues R1-R4 and the C-terminal domain are coloured purple, blue, green, yellow and orange, respectively. **d,** Schematic of the PS19 Tau filament fold. Negatively charged residues are shown in red, positively charged residues in blue, polar residues in green, non-polar residues in white, sulphur-containing residues in yellow, prolines in purple and glycines in pink. Thick connecting lines with arrowheads indicate ý-strands (ý1-ý8). The additional density is shown in red.

The four-layered ‘head’, which is made of β2-β5, consists of the nine C-terminal residues of R2 and the whole of R3. The ‘tail’ is two-layered, with one layer consisting of β1 that is made of the rest of R2 and a small portion of R1, whereas the other layer consists of β6-β8 and is formed by 28 residues of R4. Similar to filaments from line Tg2541, tau filaments from the brains of PS19 mice contain putative non-proteinaceous cofactors and/or post-translational modifications. A large external density is found next to K290 and the adjacent Q288.

### Comparison of the structures of tau filaments from Tg2541 and PS19 mice

The structures of tau filaments from Tg2541 and PS19 mice differ from each other and from the tau filament structures determined so far, regardless of their origin (extracted from brains, seeded in cells or assembled *in vitro*). However, the structures of tau filaments from mouse lines Tg2541 and PS19 share a substructure at the junction of R2 and R3, which comprises residues I297-V312. It contains mutation P301S, is stabilised by extensive hydrophobic interactions between residues and may represent a common early intermediate in filament assembly (Figure 5). Similar substructures with the wild-type P301 sequence were found previously in two other tau filament structures: one type of filament was extracted from the brain of an individual with limbic-predominant neuronal inclusion body 4R tauopathy type 1a (LNT, PDB:7P6A) [40]; the other type was assembled *in vitro* from recombinant tau (244-391) in the presence of Na2P2O7 (20a, PDB:7QL2) [41] (Figure 5a).

**Figure 5.**
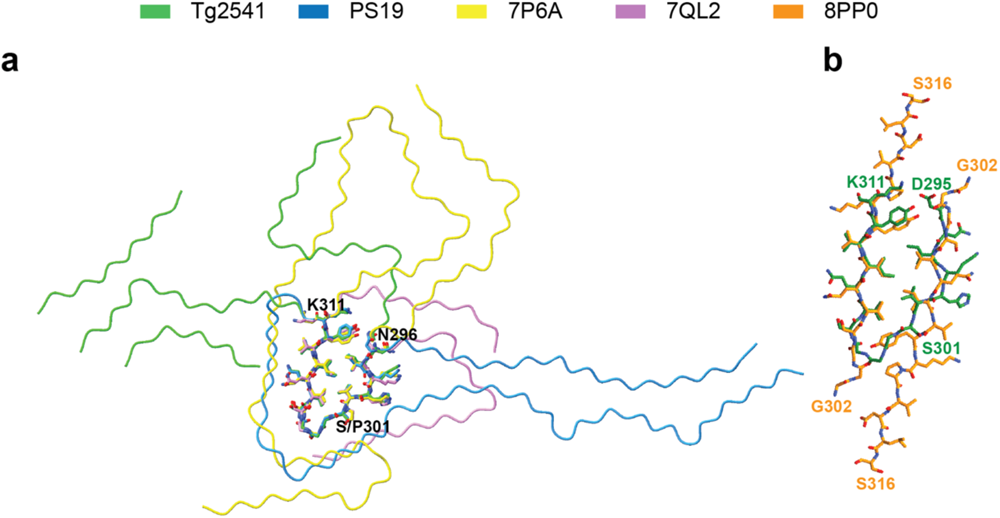
Common structural motif of Tg2541 and PS19 Tau filament folds. **a,** Superposition of Tau filament folds with a common substructure (N296-K311, sticks). P301S Tau folds from transgenic mouse lines Tg2541 and PS19 are shown in green and blue, whereas the wild-type Tau folds of LNT (PDB:7P6) and 20a (PDB:7QL2) are shown in magenta and yellow, respectively. **b,** Superposition of the Tg2541 common motif, shown in green, onto the structure of the first intermediate amyloid (FIA) (PDB:8PP0), shown in orange.

Mutation P301S tau creates an additional hydrogen bond between adjacent molecules that will further stabilise a hypothetical intermediate. The sequence of the common substructure significantly overlaps with that of the ordered core of the first intermediate amyloid (FIA) (Figure 5b), which forms during the *in vitro* assembly of recombinant tau (297-391) into either the paired helical filament Alzheimer fold or the chronic traumatic encephalopathy tau fold [42]. The FIA consists of ^302^GGGSVQIVYKPVDLS^316^ from two anti-parallel tau molecules, with a hydrophobic interface. Its ordered core (PDB:8PP0) explains the known importance of the ^306^VQIVYK^311^ (PHF6) motif for the assembly of full-length tau into filaments *in vitro* [43], in transfected cells [44] and in transgenic mice [21].

The hydrophobic core of the common substructure mimics the dimeric interface of the FIA, with V306, I308 and Y310 occupying equivalent positions in one chain, whereas I297 and V300 overlay onto V306 and I308 of the opposite chain. The additional hydrogen bond at S301 may promote the formation of a hypothetical monomeric intermediate rather than the dimeric FIA. In contrast, formation of this substructure in wild-type 4R tau filaments appears to be an exception, since most 4R tau filaments (extracted from human brains or assembled *in vitro*) adopt different conformations of the R2-R3 junction. In filaments assembled from recombinant tau (244-391), the formation of a common substructure may have been facilitated by the intramolecular disulphide bond between C291 and C322, closing the sequence of the R2-R3 junction in a short loop (20a, PDB:7QL2). The LNT filaments, which have a larger substructure in common with globular glial tauopathy (GGTI and GGTII) filaments [40], may assemble via a distinct early intermediate.

External cofactors and/or post-translational modifications appear to contribute to the differences between Tg2541 and PS19 filament folds by switching particular local conformations. In the Tg2541 fold, K294 and K298 co-ordinate a strong density on one side of the common substructure. In the PS19 fold, these lysine residues are located on different sides of the main chain, with K298 making a salt bridge with D295 that turns the preceding N-terminal residues in a different direction. K281, K317 and K321 co-ordinate the second additional density in the Tg2541 fold; it makes a right angle turn between strands β1 and β2, whereas in the PS19 fold, K281 is in the middle of the long β1 strand, where it is buried alongside D283, K343, D345 and D348, in the interface with R4 strands β6 and β7. Residues Q388 and K290 interact with a strong additional density on the other side of PS19 strand β1, whereas in the Tg2541 fold these residues are on different sides of the protein chain, with K290 making a right angle turn between strands β2 and β3.

The tau filament structure from PS19 mouse brains lends support to the proposed sequence assignments of islands A and B from the Tg2541 tau filament structure. The secondary structure of segment R4 of the PS19 fold is consistent with the β-hairpin that we propose to account for either island of the Tg2541 fold. Strand β6 corresponds exactly to the N-terminal strand of the β-hairpin and the interface with strand β1 of R2 is similar to the interface between island B and the homologous region of R3 in the Tg2541 fold. Strand β7 corresponds to the C-terminal part of the ý-hairpin and is connected to β6 by a two-residue turn, ^346^FK^347^, at the same position as the turn of the β-hairpin. At the C-terminus, β7 is shorter than the β-hairpin strand, but it can be readily extended through G355 to include the N-terminal part of β8.

## DISCUSSION

Mouse lines transgenic for human mutant P301S tau [9,10] are among the most widely used models of human tauopathies. They develop abundant filaments made of mutant 4R tau and show extensive neurodegeneration. Expression of wild-type tau in transgenic mice does not result in abundant filament formation [45–47]. However, following the injection of seeds from 5-6-month-old Tg2541 mice into the brains of mice expressing wild-type tau, filamentous tau inclusions formed [48]. Filament formation was also promoted in presymptomatic Tg2541 mice following seed injection [49]. Similarly, seeding and spreading of assembled tau have been demonstrated in PS19 mice [50]. Short tau filaments are the major species of seed-competent tau in the brains from Tg2541 mice [51]. The cryo-EM structures of tau filaments reported here are the first from mice transgenic for human tau.

Tg2541 mice develop a severe paraparesis, which is accompanied by a 50% loss of motor neurons in the lumbar spinal cord at around 20 weeks of age, with significant nerve cell loss from 12 weeks [9,21]. Here we show that a small number of tau filaments is already present in lumbar spinal cord at 5 weeks and that it increases until 25 weeks of age. Seeds of assembled tau are present from around the same time and increase in number in a reproducible manner [52]. We showed previously that tau filament formation in spinal cord precedes nerve cell dysfunction [21]. Similar findings were obtained in brainstem, with sarkosyl extraction showing the presence of tau filaments at 4 weeks of age.

The cryo-EM structures of tau filament cores from mouse line Tg2541 extend from residues K274-H329, with two islands, the sequences of which cannot be assigned unambiguously. By contrast, tau filament cores from mouse line PS19 extend from residues G271-P364. Thus, the structures of tau filaments from these lines transgenic for human P301S tau mutation are different. This may be the result of the expression of different tau isoforms, distinct genetic backgrounds and/or the use of different promoters (murine *Thy1* in [9] and murine *Prnp* in [10]).

Transgene dose also differed between both lines, in that Tg2541 mice were homozygous and PS19 mice were heterozygous. Homozygous PS19 mice do not breed [10].

It appears likely that the cellular environment of initial filament assembly determines a particular fold. Although the structures of tau filaments from humans expressing P301S tau are not known, they are probably different from those reported here, because all three 4R tau isoforms assemble and the promoter is that of *MAPT*.

The structures of tau filaments from human brains comprise R3, R4 and 10-13 amino acids after R4 [24,39,40,53,54]. By immunogold negative stain EM, they are decorated by BR133 and BR134, but not by BR136, anti-4R, BR135 or TauC4. Tau filaments from line Tg2541 comprise the C-terminal amino acid of R1, the whole of R2 and the N-terminal 24 residues of R3; they were decorated by BR133, BR134 and TauC4, but not by BR136, anti-4R or BR135. Our putative assignment of the disconnected islands with the N-terminal sequence of R4, residues 336-357 of tau, is consistent with the decoration by antibody TauC4, which was raised against residues 354-369 [26]. Tau filaments from line PS19 consist of the C-terminal four amino acids of R1, the whole of R2, the whole of R3 and the 28 N-terminal amino acids of R4. Neither filament contains the whole of R4 or the tau sequence after R4.

The lengths of tau filament cores from Tg2541 and PS19 mice are intermediate between those of filaments assembled from full-length recombinant tau in the presence of heparin [55] and filaments extracted from human brains [53]. Heparin tau filaments are decorated by BR133, BR136, TauC4 and BR134, but not by anti-4R or BR135.

Tau filaments were made of mutant human P301S tau, indicating that mouse tau did not co-assemble with human mutant tau. This is in accordance with previous studies, which showed that filament formation of human P301S tau and neurodegeneration were not affected by the presence of mouse tau [21]. It is also reminiscent of Western blotting results of sarkosyl-insoluble tau from the brains of individuals with mutation P301L tau using antibodies specific for P301L tau [56,57]. Moreover, aggregates of P301L tau could seed the assembly of P301L tau, but not of wild-type tau [58].

At end-stage, assembled tau from mice transgenic for human P301S tau is hyperphosphorylated [9,10,19]. Many of these sites were already phosphorylated in the brainstem from 2-month-old mice from line Tg2541 [57]. It remains to be seen if hyperphosphorylation is necessary for the assembly of tau into filaments. Phosphorylated sites (pT181, pS199, pS202, pT231, pS400, pT403, pS404) were located outside the core region of tau filaments. Besides phosphorylation, citrullination and methylation of the fuzzy coat of tau filaments, but not ubiquitination, were also observed in 5-month-old Tg2541 mice [59].

Filaments from mouse knock-in line *App*^NL-F^ that are made of wild-type human Aβ42 are identical to Type II Aβ42 filaments from human brains [22]. By contrast, in the presence of Arctic mutation E693G (knock-in line *App*^NL-G-F^), different structures of Aβ42 filaments were present [60,61]. These findings indicate that a missense mutation in Aβ42 is sufficient to cause a change in filament structure following expression downstream of the *App* promoter. It remains to be seen if this is also the case of tau filaments from human brains with missense mutations in the core region. In *App*^NL-G-F^ mice, identical filament structures were observed at 12 and 22 months of age. The same was observed here for Tg2541 mice aged 2 and 6 months, indicating that the filament structures did not change over time.

## Acknowledgements

This work was supported by the Electron Microscopy Facility of the MRC Laboratory of Molecular Biology. We thank Jake Grimmett, Toby Darling and Ivan Clayson for help with high-performance computing, and Sofia Lövestam for helpful discussions. We also thank the staff at ARES and the LMB Biological Services Group for their help with mouse husbandry and tissue collection. For the purpose of open access, the MRC Laboratory of Molecular Biology has applied a CC BY public copyright licence to any Author Accepted Manuscript version arising.

## Author contributions

JM and ILL characterised and maintained the Tg2541 and PS19 mouse colonies; MS, RAC and MG prepared filaments and performed immunogold negative-stain electron microscopy; MS performed cryo-EM data acquisition; MS, AGM and SHWS performed cryo-EM structure determination; SHWS and MG supervised the project and all authors contributed to the writing of the manuscript.

## Funding

This work was funded by the Medical Research Council, as part of UK Research and Innovation (MC-UP-A025-1013 to SHWS and MC-U105184291 to MG). It was also supported by Eli Lilly and Company (to MG) and the Rainwater Charitable Foundation (to MG).

## Availability of data and materials

Cryo-EM maps have been deposited in the Electron Microscopy Data Bank (EMDB) with the accession numbers EMD-18269 (Tg2541) and EMD-18268 (PS19). Corresponding refined atomic models have been deposited in the Protein Data Bank (PDB) under accession numbers PDB:8Q96 (Tg2541) and PDB:8Q92 (PS19). Please address requests for materials to the corresponding authors.

## Competing interests

The authors declare that they have no competing interests.

## SUPPLEMENTARY TABLE

### Cryo-EM Data Collection, Refinement and Validation Statistics

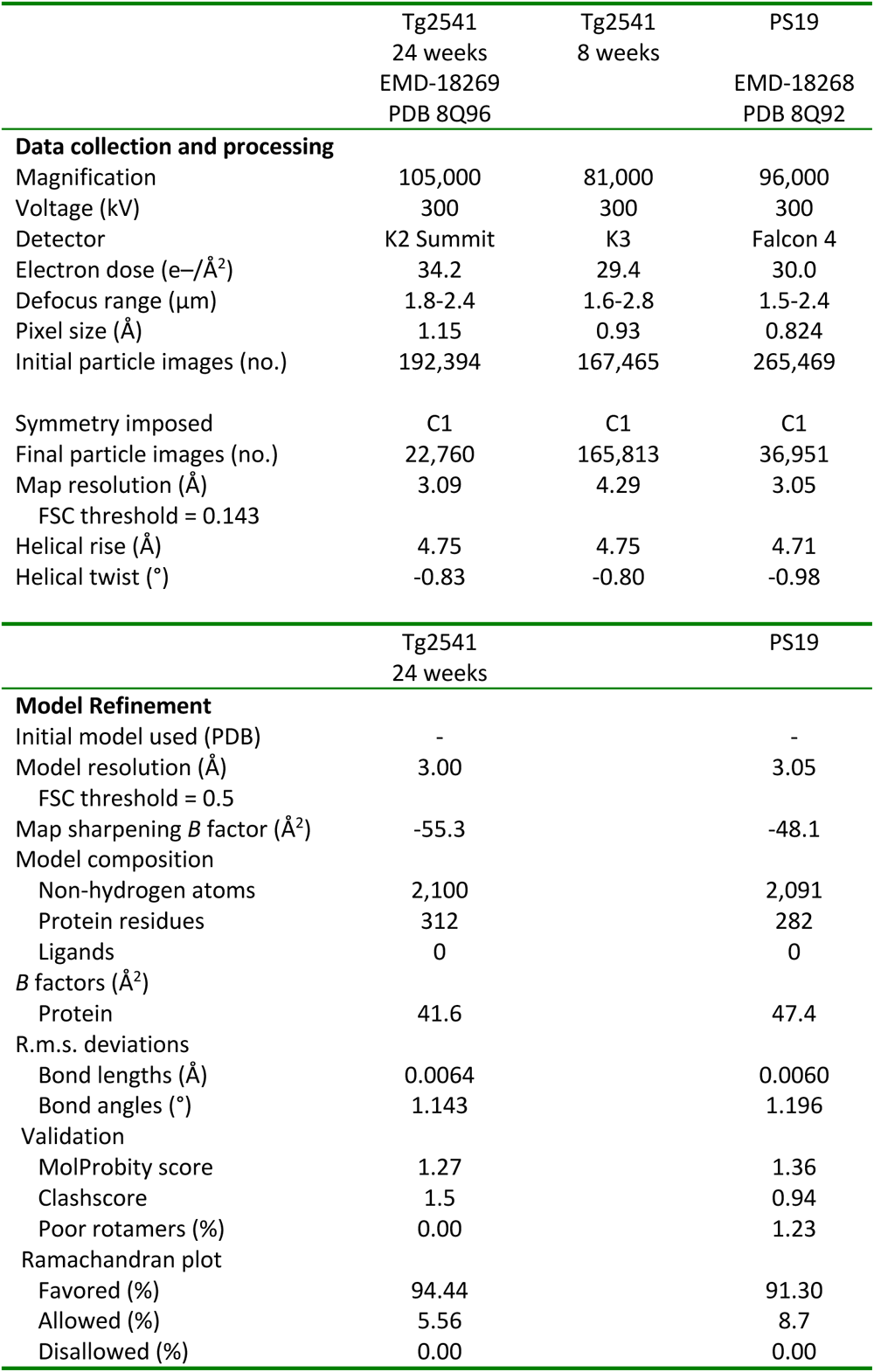

#### SUPPLEMENTARY FIGURE LEGENDS

**Supplementary Figure 1.**
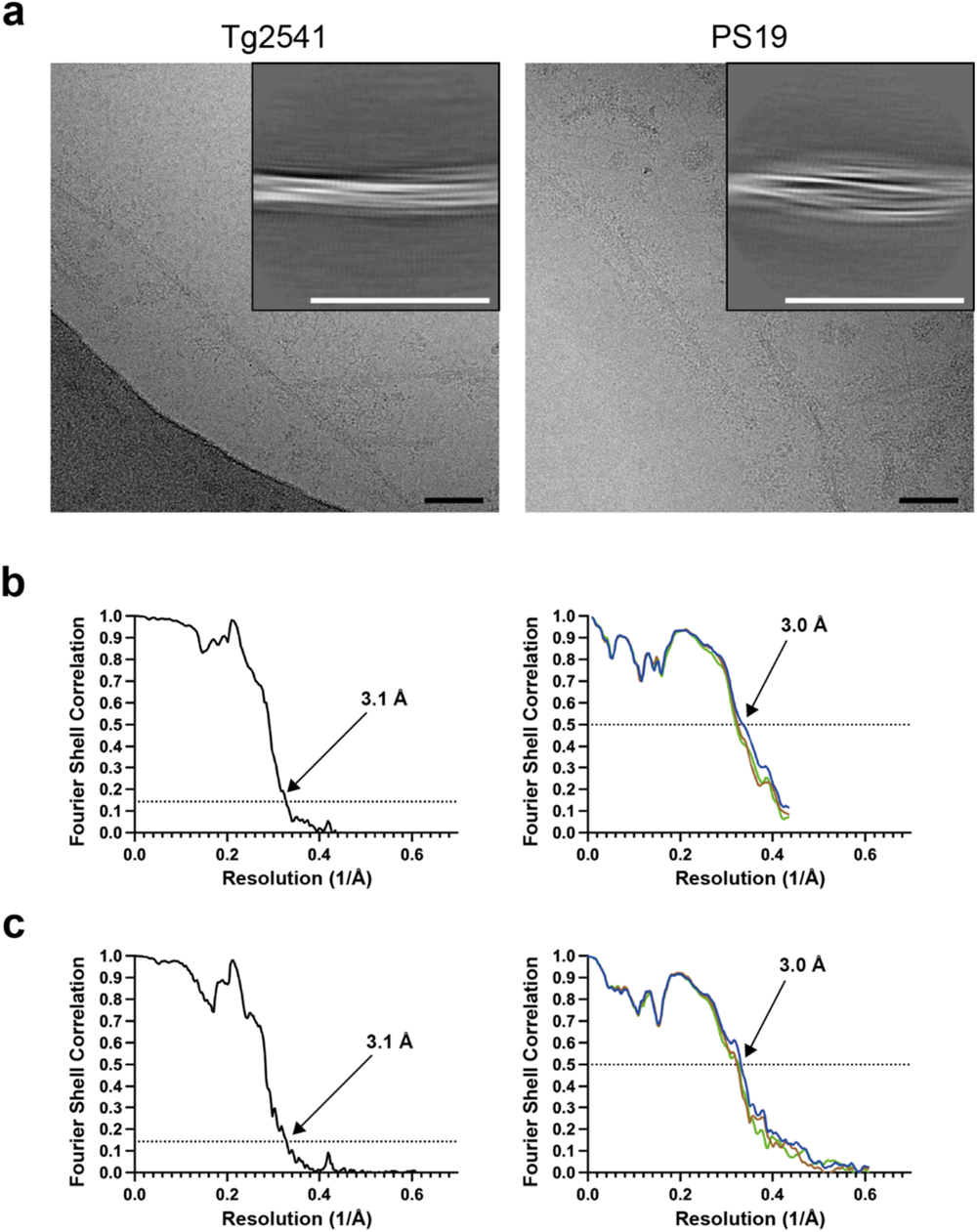
Cryo-EM 2D classifications and resolution estimates. **a,** Representative electron cryo-micrographs and 2D classification images (insets) of tau filaments from mouse lines Tg2541 and PS19 that are transgenic for human P301S tau. **b,c,** Solvent-corrected Fourier shell correlation (FSC) curves of cryo-EM half-maps (left panels) and model-to-map validation (right panels) for P301S filaments from Tg2541 (b) and PS19 mice (c). FSC curves between the model refined in the combined map versus the combined map are shown in blue; FSC curves between a model refined in half-map 1 versus half-map 1 are shown in brown (model 1 versus half-map 1); FSC curves between the same model versus half-map 2 are shown in green (model 1 versus half-map 2).

**Supplementary Figure 2.**
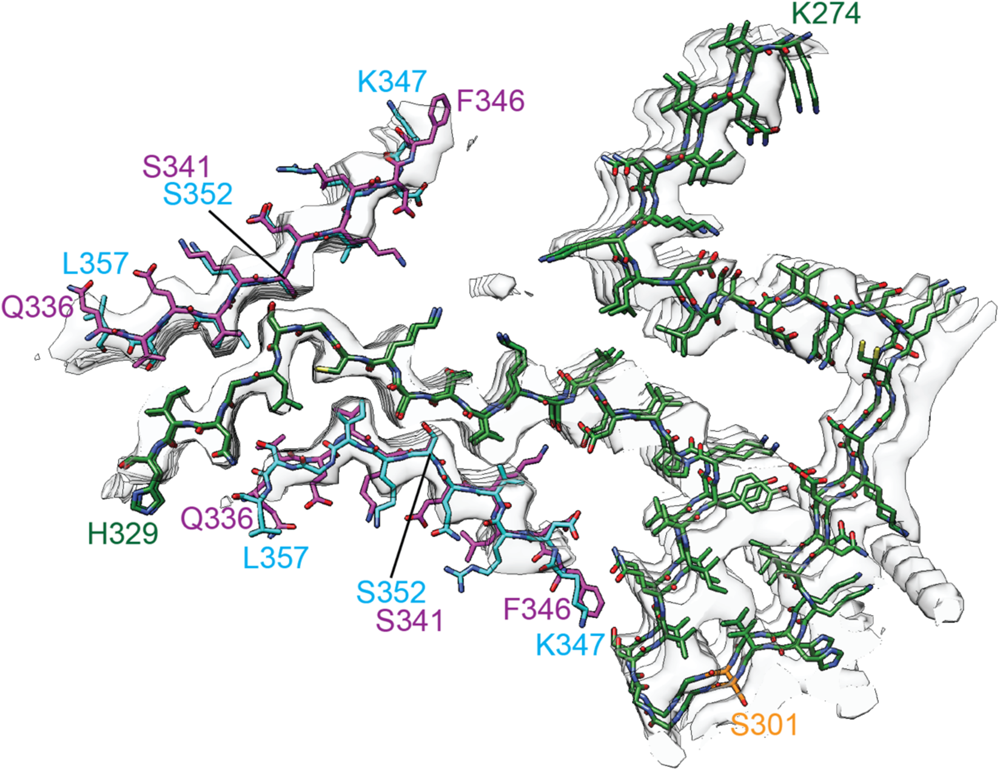
Proposed model for islands A and B of the Tg2541 Tau filament fold. The densities of both islands on two adjacent rungs were fitted with ý-hairpins comprising tau residues 336-357. N-terminal strand 336-346 is shown in magenta and C-terminal strand 347-357 in cyan. Contiguous segment 274-329 is shown in green, with the mutation site P301S shown in orange. Residues S341 and S352 that are in ‘anchor’ positions are labelled.

## Notes

### Competing Interest Statement

The authors have declared no competing interest.

## REFERENCES

1. Poorkaj P, Bird TD, Wijsman E, Nemens E, Garruto RM, Anderson L, Andreadis A, Wiederholt WC, Raskind M, Schellenberg GD (1998) Tau is a candidate gene for chromosome 17 frontotemporal dementia. Ann Neurol 43: 815–825.

2. Hutton M, Lendon CL, Rizzu P, Baker M, Froelich S, Houlden H, Pickering-Brown S, Chakraverty S, Isaacs A, Grover A et al (1998) Association of missense and 5’-splice site mutations in *tau* with the inherited dementia FTDP-17. Nature 393: 702–705.

3. Spillantini MG, Murrell JR, Goedert M, Farlow MR, Klug A, Ghetti B (1998) Mutation in the tau gene in familial multiple system tauopathy with presenile dementia. Proc Natl Acad Sci USA 95: 7737–7741.

4. Kovacs GG, Ghetti B, Goedert M (2023) Tau Proteinopathies. In: Greenfield’s Neuropathology, 10^th^ Edition (C Smith, T Jacques, GG Kovacs and A Perry, eds). CRC Press, Boca Raton, in press.

5. Goedert M, Spillantini MG, Jakes R, Rutherford D, Crowther RA (1989) Multiple isoforms of human microtubule-associated protein tau: sequences and localization in neurofibrillary tangles of Alzheimer’s disease. Neuron 3: 519–526.

6. Goedert M, Eisenberg DS, Crowther RA (2017) Propagation of tau aggregates and neurodegeneration. Annu Rev Neurosci 40: 189–210.

7. Lewis J, McGowan E, Rockwood J, Melrose H, Nacharaju P, van Slegtenhorst M, Gwinn-Hardy K, Murphy M, Baker M, Yu X, et al (2000) Neurofibrillary tangles, amyotrophy and progressive motor disturbance in mice expressing mutant (P301L) tau protein. Nature Genet 25: 402–405.

8. Götz J, Chen F, Barmettler R, Nitsch RM (2001) Tau filament formation in transgenic mice expressing P301L tau. J Biol Chem 276: 529–534.

9. Allen B, Ingram E, Takao M, Smith MJ, Jakes R, Virdee K, Yoshida H, Holzer M, Craxton M, Emson PC et al (2002) Abundant tau filaments and nonapoptotic neurodegeneration in transgenic mice expressing human P301S tau protein. J Neurosci 22: 9340–9351.

10. Yoshiyama Y, Higuchi M, Zhang B, Huang SM, Iwata N, Saido TC, Maeda J, Suhars T, Trojanowski JQ, Lee VMY (2007) Synapse loss and microglial activation precede tangles in a P301S tauopathy mouse model. Neuron 53: 553–571 (2007).

11. Bugiani O, Murrell JR, Giaccone G, Hasegawa M, Ghigo G, Tabaton M, Morbin M, Primavera A, Carella F, Solaro C et al (1999) Frontotemporal dementia and corticobasal degeneration in a family with a P301S mutation in tau. J Neuropathol Exp Neurol 58: 667–677.

12. Sperfeld AD, Collatz MB, Baier H, Palmbach M, Storch A, Schwarz J, Tatsch K, Reske S, Joosse M, Heutink P et al (1999) FTDP-17: an early-onset phenotype with parkinsonism and epileptic seizures caused by a novel mutation. Ann Neurol 46: 708–715.

13. Yasuda M, Yokoyama K, Nakayasu T, Nishimura Y, Matsui M, Yokoyama T, Miyoshi K, Tanaka C (2000) A Japanese patient with frontotemporal dementia and parkinsonism by a tau P301S mutation. Neurology 55: 1224–1227.

14. Lossos A, Reches A, Gal A, Newman JP, Soffer D, Gomori JM, Boher M, Ekstein D, Biran I, Meimner Z et al (2003) Frontotemporal dementia and parkinsonism with the P301S *tau* gene mutation in a Jewish family. J Neurol 250: 733–740.

15. Werber E, Klein C, Grünfeld J, Rabey JM (2003) Phenotypic presentation of frontotemporal dementia with parkinsonism-chromosome 17 type P301S in a patient of Jewish-Algerian origin. Mov Disord 18: 595–598.

16. Yasuda M, Nakamura Y, Kawamata T, Kaneyuki H, Maeda K, Komure O (2005) Phenotypic heterogeneity within a new family with the *MAPT* P301S mutation. Ann Neurol 58: 920–928.

17. Goedert M, Jakes R, Crowther RA (1999) Effects of frontotemporal dementia FTDP-17 mutations on heparin-induced assembly of tau filaments. FEBS Lett 450: 306–311.

18. D’Souza I, Poorkaj P, Hong M, Nochlin D, Lee VMY, Bird TD, Schellenberg GD (1999) Missense and silent tau gene mutations cause frontotemporal dementia with parkinsonism – chromosome 17 type, by affecting multiple alternative RNA splicing regulatory elements. Proc Natl Acad Sci USA 96: 5598–5603.

19. Delobel P, Lavenir I, Fraser G, Ingram E, Holzer M, Ghetti B, Spillantini MG, Crowther RA, Goedert M (2008) Analysis of tau phosphorylation and triuncation in a mouse model of human tauopathy.

20. Velasco A, Fraser G, Delobel P, Ghetti B, Lavenir I, Goedert M (2008) Detection of filamerntous tau inclusions by the fluorescent Congo red derivative [(*trans, trans*)-1-fluoro-2,5-bis(3-hydroxycarbonyl-4-hydroxy)styrylbenzene]. FEBS Lett 582: 901–906 (2008).

21. Macdonald JA, Bronner IF, Drynan L, Fan J, Curry A, Fraser G, Lavenir I, Goedert M (2019) Assembly of transgenic human P301S Tau is necessary for neurodegeneration in murine spinal cord. Acta Neuropathologica Commun 7: 44.

22. Yang Y, Arseni D, Zhang W, Huang M, Lövestam S, Schweighauser M, Kotecha A, Murzin AG, Peak-Chew SY, Macdonald J et al (2022) Cryo-EM structures of amyloid-β 42 filaments from human brains. Science 375: 167–172.

23. Goedert M, Spillantini MG, Cairns NJ, Crowther RA (1992) Tau proteins of Alzheimer paired helical filaments: abnormal phosphorylation of all six brain isoforms. Neuron 8: 159–168.

24. Falcon B, Zhang W, Murzin AG, Murshudov G, Garringer HJ, Vidal R, Crowther RA, Ghetti B, Scheres SHW, Goedert M (2018) Structures of filaments from Pick’s disease reveal a novel tau protein fold. Nature 561: 137–140.

25. Dan A, Takahashi M, Masuda-Suzukake M, Kametani F, Nonaka T, Kondo H, Akiyama H, Arai T, Mann DMA, Saito Y et al (2013) Extensive deamidation at asparagine residue 279 accounts for weak immunoreactivity of tau with RD4 antibody in Alzheimer’s disease brain. Acta Neuropathol Commun 1: 54.

26. Taniguchi-Watanabe S, Arai T, Kametani F, Nonaka T, Masuda-Suzukake M, Tarutani A, Murayama S, Saito Y, Arima K, Yoshida M et al (2016) Biochemical classification of tauopathies by immunoblot, protein sequence and mass spectrometric analyses of sarkosyl-insoluble and trypsin-resistant tau. Acta Neuropathol 131: 267–280.

27. He S, Scheres SHW (2017) Helical reconstruction in RELION. J Struct Biol 193: 163–176.

28. Zivanov J, Nakane T, Forsberg BO, Kimanius D, Hagen WJ, Lindahl E, Scheres SHW (2018) New tools for automated high-resolution cryo-EM structure determination in RELION-3. eLife 7: e42166.

29. Rohou A, Grigorieff N (2015) CTFFIND4: fast and accurate defocus estimation from electron micrographs. J Struct Biol 192: 216–221.

30. Zivanov J, Otón J, Ke Z, von Kügelen A, Pyle E, Qu K, Morado D, Castaño-Diez D, Zanetti G, Bharat TAM, et al (2022) A Bayesian approach to single-particle electron cryo-tomography in RELION-4.0. eLife 11: e83724.

31. Scheres SHW (2020) Amyloid structure determination in RELION-3.1. Acta Cryst D 76: 94–101.

32. Emsley P, Lohkamp B, Scott WG, Cowtan K (2010) Features and development of Coot. Acta Crystallogr D 66: 486–501.

33. Yamashita K, Palmer CM, Burnley T, Murshudov GN (2021) Cryo-EM single-particle structure refinement and map calculation using *Servalcat*. Acta Crystallogr D 53: 240–255.

34. Murshudov GN, Vagin AA, Dodson EJ (1997) Refinement of macromolecular structures by the maximum-likelihood method. Acta Crystallogr D 53: 240–255.

35. Murshudov GN, Skubák P, Lebedev AA, Pannu NS, Steiner RA, Nicholls RA, Winn MD, Long F, Vagin AA (2011) REFMAC5 for the refinement of macromolecular crystal structures. Acta Crystallogr D 67: 355–367.

36. Chen VB, Arendall WB, Headd JJ, Keedy DA, Immormino RM, Kaprai GJ, Murray LW, Richardson JS, Richardson DC (2010) MolProbity: all-atom structure validation for macromolecular crystallography. Acta Crystallogr D 66: 12–21.

37. Pettersen EF, Goddard TD, Huang CC, Meng EC, Couch GS, Croll TI, Morris JH, Ferrin TE (2021) ChimeraX: structure visualization for researchers, educators, and developers. Protein Sci 30: 70–82.

38. Schrödinger L, DeLano W (2020) PyMOL available at: <http://www.pymol.org/pymol>.

39. Zhang W, Tarutani A, Newell KL, Murzin AG, Matsubara T, Falcon B, Vidal R, Garringer HJ, Shi Y, Ikeuchi T et al (2020) Novel tau filament fold in corticobasal degeneration. Nature 580: 283–287.

40. Shi Y, Zhang W, Yang Y, Murzin AG, Falcon B, Kotecha A, van Beers M, Tarutani A, Kametani F, Garringer HJ, et al (2021) Structure-based classification of tauopathies. Nature 598: 359–363.

41. Lövestam S, Koh, FA, van Knippenberg B, Kotecha A, Murzin AG, Goedert M, Scheres SHW (2022) Assembly of recombinant tau into filaments identical to those of Alzheimer’s disease and chronic traumatic encephalopathy. eLife 11: e76494.

42. Lövestam S, Li D, Wagstaff JL, Kotecha A, Kimanius D, McLaughlin SH, Murzin AG, Freund SMV, Goedert M, Scheres SHW (2023) Disease-specific tau filaments assemble via polymorphic intermediates. BioRxiv.

43. Von Bergen M, Friedhoff P, Biernat J, Heberle J, Mandelkow EM, Mandelkow E (2000) Assembly of tau protein into Alzheimer paired helical filaments depends on a local sequence motif (^306^VQIVYK^311^) forming β structure. Proc Natl Acad Sci USA 97: 5129–5134.

44. Falcon B, Cavallini A, Angers R, Glover S, Murray TK, Barnham L, Jackson S, O’Neill MJ, Isaacs AM, Hutton ML et al (2015) Conformation determines the seeding potencies of native and recombinant tau aggregates. J Biol Chem 290: 1049–1065.

45. Götz J, Probst A, Spillantini MG, Schäfer T, Jakes R, Bürki K, Goedert M (1995) Somatodendritic localisation and hyperphosphorylation of tau protein in transgenic mice expressing the longest human brain tau isoform. EMBO J 14: 1304–1313.

46. Probst A, Götz J, Wiederhold KH, Tolnay M, Mistl C, Jaton AL, Hong M, Ishihara T, Lee VMY, Trojanowski JQ et al (2000) Axonopathy and amyotrophy in mice transgenic for human four-repeat tau protein. Acta Neuropathol 99: 469–481.

47. Sahara N, Yanai R (2023) Limitations of human tau-expressing mouse models and novel approaches of mouse modelling for tauopathy. Front Neurosci 17: 11349761.

48. Clavaguera F, Bolmont T, Crowther RA, Abramowski D, Frank S, Probst A, Fraser G, Stalder AK, Beibel M, Staufenbiel M et al (2009) Transmission and spreading of tauopathy in transgenic mouse brain. Nature Cell Biol 11: 909–913.

49. Ahmed Z, Cooper J, Murray TK, Garn K, McNaughton E, Clarke H, Parhizkar S, Ward MA, Cavallini A, Jackson S et al (2014) A novel *in vivo* model of tau propagation with rapid and progressive neurofibrillary tangle pathology: the pattern of spread is determined by connectivity, not proximity. Acta Neuropathol 127: 667–683.

50. Boluda S, Iba M, Zhang B, Raible KM, Lee VMY, Trojanowski JQ (2015) Differential induction and spread of tau pathology in young PS19 tau transgenic mice following intracerebral injections of pathological tau from Alzheimer’s disease or corticobasal degeneration brains. Acta Neuropathol 129: 221–237.

51. Jackson SJ, Kerridge C, Cooper J, Cavallini A, Falcon B, Cella CV, Landi A, Szekeres PG, Murray TK, Ahmed Z et al (2016) Short fibrils constitute the major species of seed-competent tau in the brains of mice transgenic for human P301S tau. J Neurosci 36: 762–772.

52. Woerman AL, Patel S, Kazmi SA, Oehler A, Freyman Y, Espiritu L, Cotter R, Castaneda JA, Olson SH, Prusiner SB (2017) Kinetics of human mutant tau prion formation in the brains of 2 transgenic mouse lines. JAMA Neurol 74: 1464–1472.

53. Fitzpatrick AWP, Falcon B, He S, Murzin AG, Murshudov G, Garringer HJ, Crowther RA, Ghetti B, Goedert M (2017) Cryo-EM structures of tau filaments from Alzheimer’s disease. Nature 547: 185–190.

54. Falcon B, Zivanov J, Zhang W, Murzin AG, Garringer HJ, Vidal R, Crowther RA, Newell KL, Ghetti B, Goedert M et al (2019) Novel tau filament fold in chronic traumatic encephalopathy encloses hydrophobic molecules. Nature 568: 420–423.

55. Zhang W, Falcon B, Murzin AG, Fan J, Crowther RA, Goedert M, Scheres SHW (2019) Heparin-induced tau filaments are polymorphic and differ from those in Alzheimer’s and Pick’s diseases. eLife 8: e43584.

56. Rizzu P, Joosse M, Ravid R, Hoogeveen A, Kamphorst W, van Swieten JC, Willemsen R, Heutink P (2000) Mutation-dependent aggregation of tau protein and its selective depletion from the soluble fraction in brain of P30L FTDP-17 patients. Hum Mol Genet 9: 3075–3082.

57. Miyasaka T, Morishima-Kawashima M, Ravid R, Kamphorst W, Nagashima K, Ihara Y (2001) Selective deposition of mutant tau in the FTDP-17 brain affected by the P301L mutation. J Neuropathol Exp Neurol 60: 872–884.

58. Aoyagi H, Hasegawa M, Tamaoka A (2007) Fibrillogenic nuclei composed of P301L mutant tau induce elongation of P301L tau, but not wild-type tau. J Biol Chem 282: 20309–20318.

59. Wenger K, Viode A, Schlaffner CN, van Zalm P, Cheng L, Dellovade T, Langlois X, Bannon A, Chang R, Connors TR, et al (2023) Common mouse models of tauopathy reflect early but not late human disease. Mol Neurodegen 18: 10.

60. Yang Y, Zhang W, Murzin AG, Schweighauser M, Huang M, Lövestam S, Peak-Chew SY, Saito T, Saido TC, Macdonald J et al (2023) Cryo-EM structures of amyloid-β filaments with the Arctic mutation (E22G) from human and mouse brains. Acta Neuropathol 145: 325–333.

61. Leistner C, Wilkinson M, Burgess A, Lovatt M, Goodbody S, Xu Y, Deuchars S, Radford SE, Ranson NA, Frank RAW (2023) The in-tissue molecular architecture of β-amyloid pathology in the mammalian brain. Nature Commun 14: 2833.

